# Reovirus infection is regulated by NPC1 and endosomal cholesterol homeostasis

**DOI:** 10.1101/2021.09.27.462002

**Authors:** Paula Ortega-Gonzalez, Gwen Taylor, Rohit K. Jangra, Raquel Tenorio, Isabel Fernández de Castro, Bernardo A. Mainou, Robert C. Orchard, Craig B. Wilen, Pamela H. Brigleb, Jorna Sojati, Kartik Chandran, Cristina Risco, Terence S. Dermody

## Abstract

Cholesterol homeostasis is required for the replication of many viruses, including Ebola virus, hepatitis C virus, and human immunodeficiency virus-1. Niemann-Pick C1 (NPC1) is an endosomal-lysosomal membrane protein involved in cholesterol trafficking from late endosomes and lysosomes to the endoplasmic reticulum. We identified NPC1 in CRISPR and RNA interference screens as a putative host factor for infection by mammalian orthoreovirus (reovirus). Following internalization via clathrin-mediated endocytosis, the reovirus outer capsid is proteolytically removed, the endosomal membrane is disrupted, and the viral core is released into the cytoplasm where viral transcription, genome replication, and assembly take place. We found that reovirus infection is significantly impaired in cells lacking NPC1, but infection is restored by treatment of cells with hydroxypropyl-β-cyclodextrin, which binds and solubilizes cholesterol. Absence of NPC1 did not dampen infection by infectious subvirion particles, which are reovirus disassembly intermediates that bypass the endocytic pathway for infection of target cells. NPC1 is not required for reovirus attachment to the plasma membrane, internalization into cells, or uncoating within endosomes. Instead, NPC1 is required for delivery of transcriptionally active reovirus core particles into the cytoplasm. These findings suggest that cholesterol homeostasis, ensured by NPC1 transport activity, is required for reovirus penetration into the cytoplasm, pointing to a new function for NPC1 and cholesterol homeostasis in viral infection.

**Author summary:** Genetic screens are useful strategies to identify host factors required for viral infection. NPC1 was identified in independent CRISPR and RNA interference screens as a putative host factor required for reovirus replication. We discovered that NPC1-mediated cholesterol transport is dispensable for reovirus attachment, internalization, and disassembly but required for penetration of the viral disassembly intermediate from late endosomes into the cytoplasm. These findings pinpoint an essential function for cholesterol in the entry of reovirus and raise the possibility that cholesterol homeostasis regulates the entry of other viruses that penetrate late endosomes to initiate replication.

## INTRODUCTION

Viral replication is dependent on cellular proteins and pathways for entry, transport, and release of the viral genome to sites of replication in the cell. Viral attachment to host cells occurs by interactions with cell-surface proteins, lipids, and carbohydrate moieties at the plasma membrane and often triggers virus uptake by receptor-mediated endocytosis (1–7). Viruses that traverse through endosomes must escape the endosomal compartment and release their genomes at sites of replication to initiate productive infection. Enveloped viruses generally accomplish endosomal escape using mechanisms involving receptor- or pH-mediated fusion of the viral envelope and endosomal membrane (6, 8–10). In contrast, nonenveloped viruses penetrate endosomal membranes by establishing small membrane pores or large membrane disruptions (9, 11–13). While both enveloped and nonenveloped viruses depend on conformational changes of viral structural proteins to escape endosomes, mechanisms underlying nonenveloped virus membrane penetration are not well understood (6).

Mammalian orthoreoviruses (reoviruses) are nonenveloped icosahedral viruses that infect a broad range of mammalian hosts. Reovirus infections are usually asymptomatic in humans, but these viruses have been implicated in development of celiac disease (14). Reovirus virions include two protein shells, the outer capsid, composed primarily of μ1-σ3 heterohexamers, and core (15–17). The core contains 10 segments of double-stranded (ds) RNA, which are classified by size into three large (L), three medium (M), and four small (S) segments (17). Following receptor-mediated endocytosis, the reovirus outer capsid undergoes a series of conformational changes and disassembly events required for release of transcriptionally active cores into the cytoplasm (18, 19).

Within late endosomes, acid-dependent cathepsin proteases catalyze proteolysis of the viral outer-capsid protein σ3 and cleavage of the membrane-penetration protein μ1 to δ and φ, resulting in formation of metastable intermediates termed infectious subvirion particles (ISVPs) (20–24). Endosomal lipid composition induces ISVPs to undergo additional conformational changes resulting in exposure of hydrophobic domains of δ, release of pore-forming fragment μ1N, and formation of ISVP*s (25, 26). Release of μ1N during ISVP-to-ISVP* conversion leads to endosomal penetration and liberation of the viral core into the cytoplasm where infection progresses (27–31). Although some essential viral and host factors required for reovirus penetration of the endosome are known, the process is still not well understood.

In this study, we used CRISPR and RNA interference screens to discover that Niemann Pick C1 (NPC1), an endolysosomal transmembrane protein that mediates cholesterol egress from late endosomes for redistribution to cellular membranes (32–34), is required for reovirus infection. We found that genetic ablation of NPC1 in human brain microvascular endothelial cells (HBMECs) diminishes reovirus infection by virions but not by ISVPs, suggesting that NPC1 is required for steps that differ between virions and ISVPs. Treatment of NPC1-null HBMECs with hydroxypropyl-beta-cyclodextrin (HβCD), a macrocycle that binds and solubilizes cholesterol, restored infectivity by reovirus virions, suggesting that endosomal cholesterol homeostasis contributes to efficient reovirus entry. While NPC1 is not required for viral attachment to the plasma membrane, internalization, or uncoating within endosomes, we found that NPC1 is required for efficient release of reovirus cores from endosomes into the cytoplasm. Together, these findings suggest that cholesterol homeostasis, mediated by NPC1 cholesterol transport activity, is essential for reovirus cell entry and penetration into the cytoplasm.

## RESULTS

### CRISPR/Cas-9 and siRNA screens for host factors required for reovirus infection identify NPC1

To discover host factors required for reovirus infection, we conducted genome-wide CRISPR/Cas-9 and siRNA-based cell-survival screens. The CRISPR/Cas-9 screen was conducted using BV2 mouse microglial cells with the murine Asiago sgRNA library targeting over 20,000 genes. BV2 CRISPR cell libraries were infected with reovirus strains type 1 Lang (T1L) and type 3 Dearing (T3D) and cultured for nine days prior to isolation of genomic DNA (gDNA) from surviving cells and deep sequencing. STARS analysis was conducted to identify enriched CRISPR gRNAs within the surviving cell population (Fig. 1A and Table S1). The siRNA screen was conducted using HeLa S3 cells transfected with the ON-TARGET plus siRNA whole genome library targeting over 18,000 genes (35). Transfected cells were infected with reovirus strain T3SA+ and scored for viability using an ATP-dependent luminescence assay. T3SA+ contains nine genes from T1L and the S1 gene from strain T3C44-MA (36). T3SA+ binds all known reovirus receptors and is cytolytic. Robust Z scores (median absolute deviation) were calculated for each sample (Fig. 1B and Table S2).

**Fig. 1.**
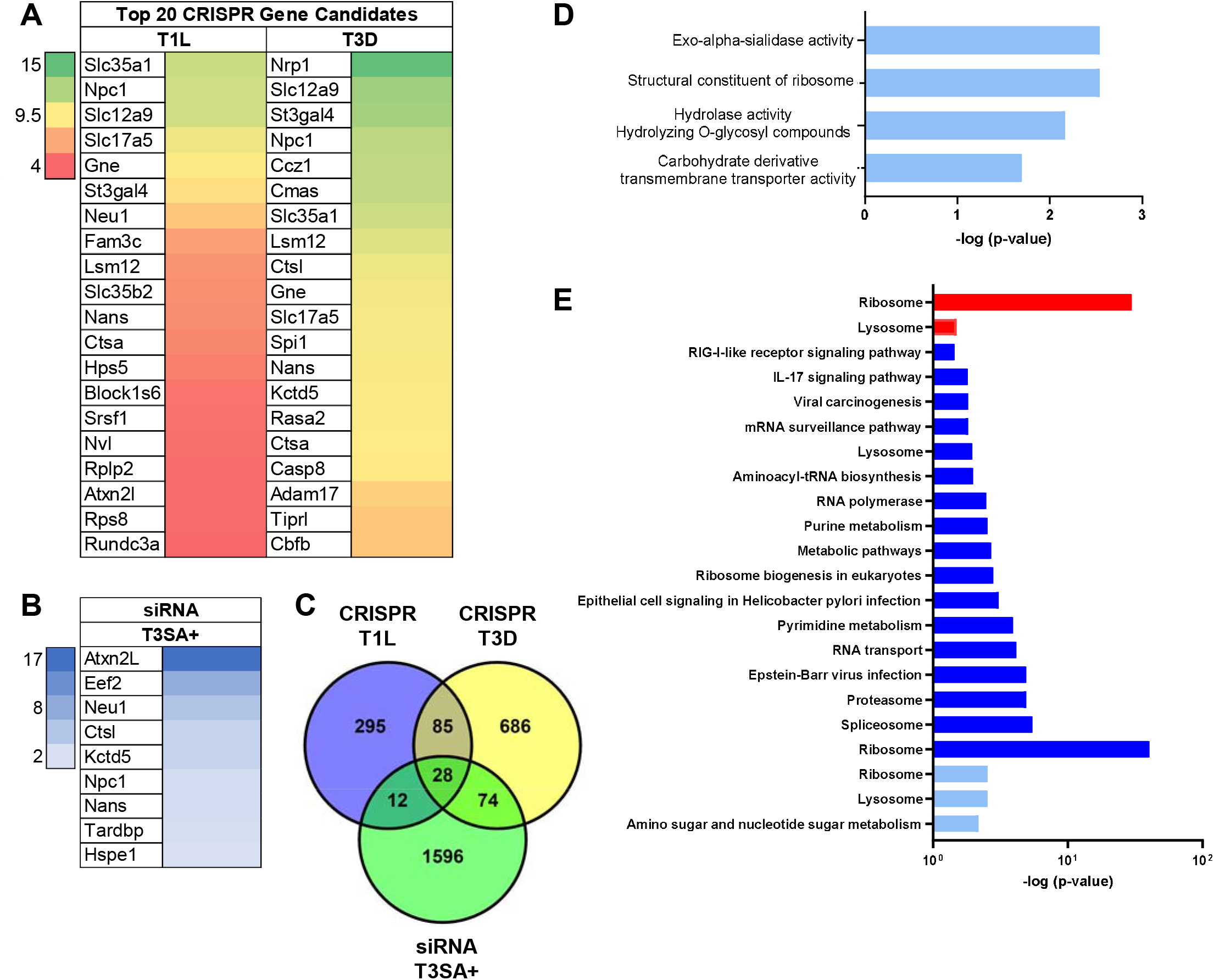
CRISPR and siRNA screens identify NPC1 as a cellular factor required for reovirus infection. (A) The top 20 candidates from the CRISPR screen using reovirus strains T1L and T3D are ranked by their STAR scores. Heat map indicates STAR values. (B) Genes from the siRNA screen using reovirus strain T3SA+ common to the CRISPR screen using T1L and T3D, excluding ribosomal genes. Heat map indicates z-score values. (C) Venn diagram of genes from the CRISPR screens using T1L and T3D and the siRNA screen using T3SA+. (D) Molecular function pathways using Gene Ontology to analyze genes from the CRISPR screen common to T1L and T3D. (E) KEGG pathways identified for the CRISPR screen using T1L (red) and T3D (blue) and siRNA screen using T3SA+ (light blue).

Key genes and pathways essential for reovirus replication were defined by comparing the CRISPR/Cas-9 and siRNA screen lists using STRING-db (Fig. 1C). In the CRISPR/Cas-9 screen, four functional pathways defined by Gene Ontology (GO) terms were common to both T1L and T3D, including sialic acid biosynthesis and metabolism (Fig. 1D). Sialic acid is a reovirus attachment factor, and genes involved in sialic acid biosynthesis and metabolism, including *Slc35a1*, are required for T3SA+ replication in BV2 cells (37). These data provide confidence that the target genes identified in the CRISPR/Cas-9 screen represent biologically significant candidates. We also compared KEGG pathways identified in the CRISPR/Cas-9 and siRNA screens to increase the likelihood of significant gene targets. Ribosome and lysosome pathways were the only pathways common to both screens (Fig. 1E). Lysosomal genes include *Ctsl*, *Neu1*, and *Npc1*. *Ctsl* encodes cathepsin L, which is required for cleavage of the reovirus outer capsid to form ISVPs (22). *Neu1* encodes neuraminidase, a lysosomal sialidase that cleaves sialic acid linkages required for reovirus infectivity (38). *Npc1* encodes NPC1, a cholesterol transporter that resides in the limiting membrane of endosomes and lysosomes (33, 34).

### Engineering and characterization of HBMECs with CRISPR-targeted Npc1

Based on the function of NPC1 in cell entry and replication of other viruses (39) and its identification in both CRISPR and siRNA screens, we evaluated a potential role for NPC1 in reovirus replication. Human brain microvascular endothelial cells (HBMECs) are susceptible to reovirus infection (40) and amenable to CRISPR/Cas-9 gene editing (41). To facilitate these studies, we used CRISPR/Cas-9 gene editing to engineer a clonal HBMEC cell line lacking the *NPC1* gene (KO cells). The NPC1 KO cells were complemented by stable transfection of a functional NPC1 allele (KO+ cells).

The newly engineered NPC1 KO and KO+ cell lines were characterized for NPC1 expression and cholesterol distribution relative to wild-type (WT) HBMECs. Expression of NPC1 in WT, KO, and KO+ cells was tested using immunoblotting. As anticipated, NPC1 expression in KO cells was abrogated relative to WT and KO+ cells (Fig. S1A). There was an observable increase in NPC1 expression in KO+ cells compared with WT cells (Fig. S1B), but the difference was not statistically significant. In the absence of functional NPC1, cholesterol reorganizes from a homogeneous distribution to accumulate in endosomal compartments (32, 33). To define the distribution of cholesterol in NPC1-null HBMECs, we used fluorescent filipin III to label cholesterol in fixed cells and imaged cholesterol distribution using fluorescence microscopy (Fig. S1C). Cholesterol distribution was homogeneous in WT (Fig. S1C, left) and KO+ cells (Fig. S1C, right). However, cholesterol accumulated around the nucleus in KO cells (Fig. S1C, center) in a pattern consistent with the distribution of endosomes (Fig. S1D), confirming the absence of functional NPC1. Thus, KO cells display the expected phenotype of altered cholesterol distribution when NPC1-dependent cholesterol transport is disrupted. Furthermore, complementing NPC1 expression in KO cells restores the normal distribution of cholesterol, demonstrating that the observed phenotype is specific for NPC1 expression.

### Reovirus infection by virions but not by ISVPs is impaired in NPC1 KO cells

ISVPs prepared by treatment of virions *in vitro* with intestinal or endosomal proteases bind to reovirus receptors and enter target cells by direct penetration of the plasma membrane and bypass requirements for internalization into the endocytic compartment and acid-dependent proteolysis (21, 22, 42). To determine whether NPC1 is required for reovirus replication, and further whether NPC1 mediates a step in the infectious cycle that differs between virions and ISVPs, we adsorbed WT, KO, and KO+ cells with reovirus strain T1L M1 P208S virions or ISVPs. Reovirus T1L M1-P208S contains a point mutation in the M1 gene that causes viral factories to have a globular morphology similar to the morphology of factories formed by reovirus T3D (43), which renders infected cells easier to detect. Infected cells were visualized by immunofluorescence (IF) staining for reovirus antigen at 18 h post-adsorption (Fig. 2). Following adsorption with reovirus virions, the number of infected KO cells was reduced by approximately 50% relative to infected WT and KO+ cells (Fig. 2A). A similar reduction in the number of infected KO cells relative to WT and KO+ cells was observed when WT, KO, and KO+ cells were adsorbed with T1L, T3D, and T3SA+ virions, the reovirus strains used in the CRISPR/Cas9 and siRNA screens (Fig. S2). In contrast, no significant differences in numbers of infected cells were observed following adsorption of WT, KO, and KO+ cells with ISVPs (Fig. 2B). Viral progeny production and release was determined by quantifying viral titers in cell lysates and supernatants at 0, 24, and 48 h following adsorption of WT, KO, and KO+ cells with virions or ISVPs. Following infection by virions, viral titers in lysates and supernatants of KO cells were 10- to 100-fold less than those in WT and KO+ cells (Fig. 2C and E). In contrast, following infection by ISVPs, viral titers in lysates and supernatants of all three cell types were comparable (Fig. 2D and F). Together, these results suggest that NPC1 is required for reovirus infection and functions at a step in the infectious cycle that differs between virions and ISVPs.

**Fig. 2.**
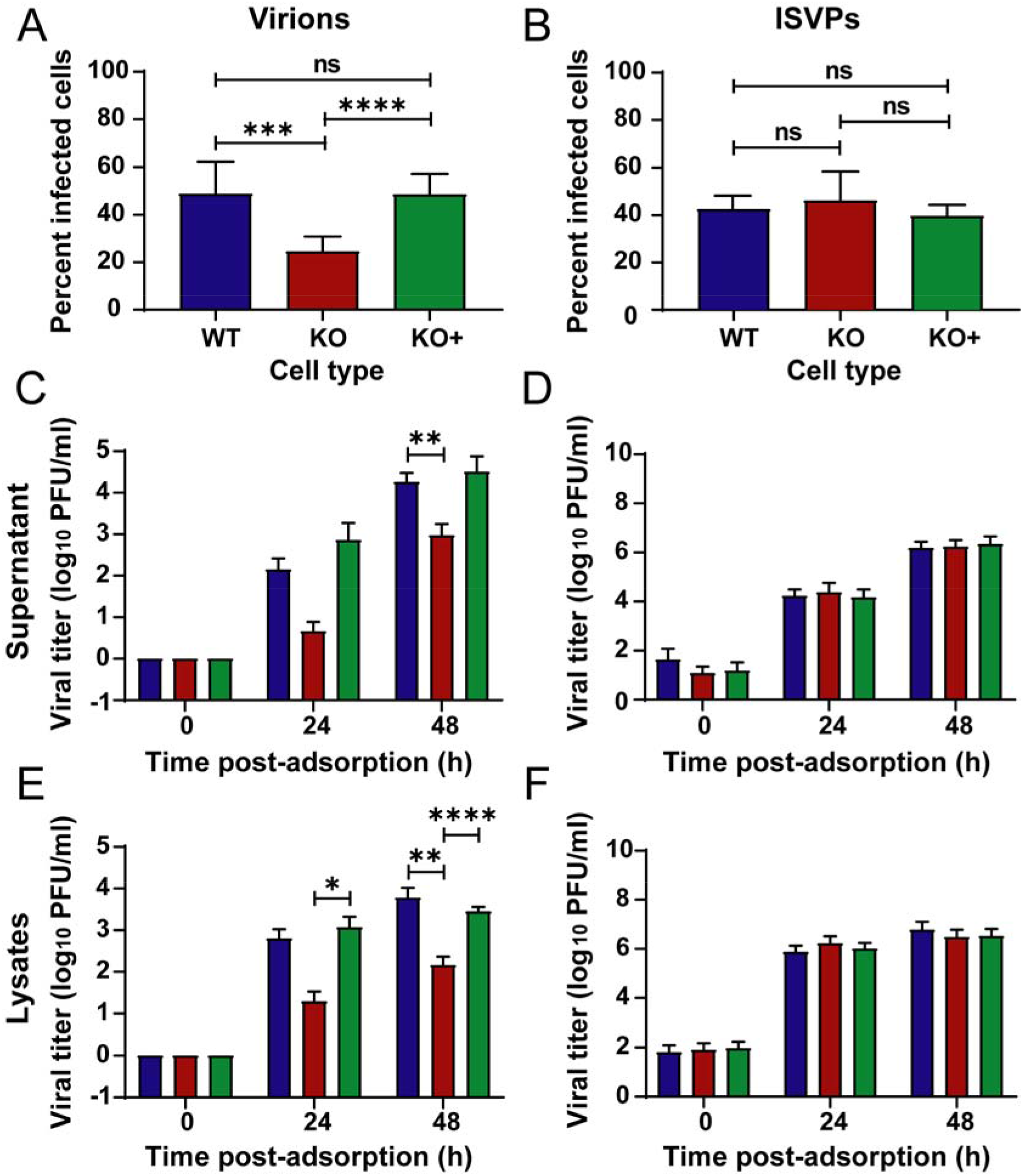
Viral infectivity and titers following adsorption by reovirus virions and ISVPs. (A, B) WT, KO, and KO+ HBMECs were adsorbed with reovirus (A) virions or (B) ISVPs at MOIs of 10,000 or 100 particles/cell, respectively, and fixed at 18 h post-adsorption. The percentage of infected cells was determined by enumerating reovirus-infected cells following immunostaining with a reovirus-specific antiserum. (C-F) WT, KO, and KO+ cells were adsorbed with reovirus (C, E) virions at an MOI of 1 PFU/cell or (D, F) ISVPs at an MOI of 5 particles/cell. Viral titers in cell-culture supernatants and lysates were determined by plaque assay at 0, 24, and 48 h post-adsorption. The results are presented as the mean of three independent experiments. Error bars indicated standard deviation. *, *P* < 0.05; **, *P* < 0.01; ***, *P* < 0.001; ****, *P* < 0.0001, as determined by t-test.

### NPC1 is not required for reovirus attachment, internalization, or uncoating

Reovirus entry can be divided into four main stages: viral binding to cell-surface receptors, viral internalization by endocytosis, proteolytic removal of the viral outer capsid, and penetration of the core from late endosomes into the cytosol (19). We characterized NPC1 KO cells for the capacity to support each step of the reovirus entry pathway to define the function of NPC1 in reovirus infection. To determine whether NPC1 is required for reovirus attachment to target cells, we quantified viral binding using flow cytometry. The quantity of virus bound to the surface of all three cell types was comparable, and no statistically significant differences were observed (Fig. 3A). These data suggest that reovirus attachment to cells is not dependent on expression of NPC1.

**Fig. 3.**
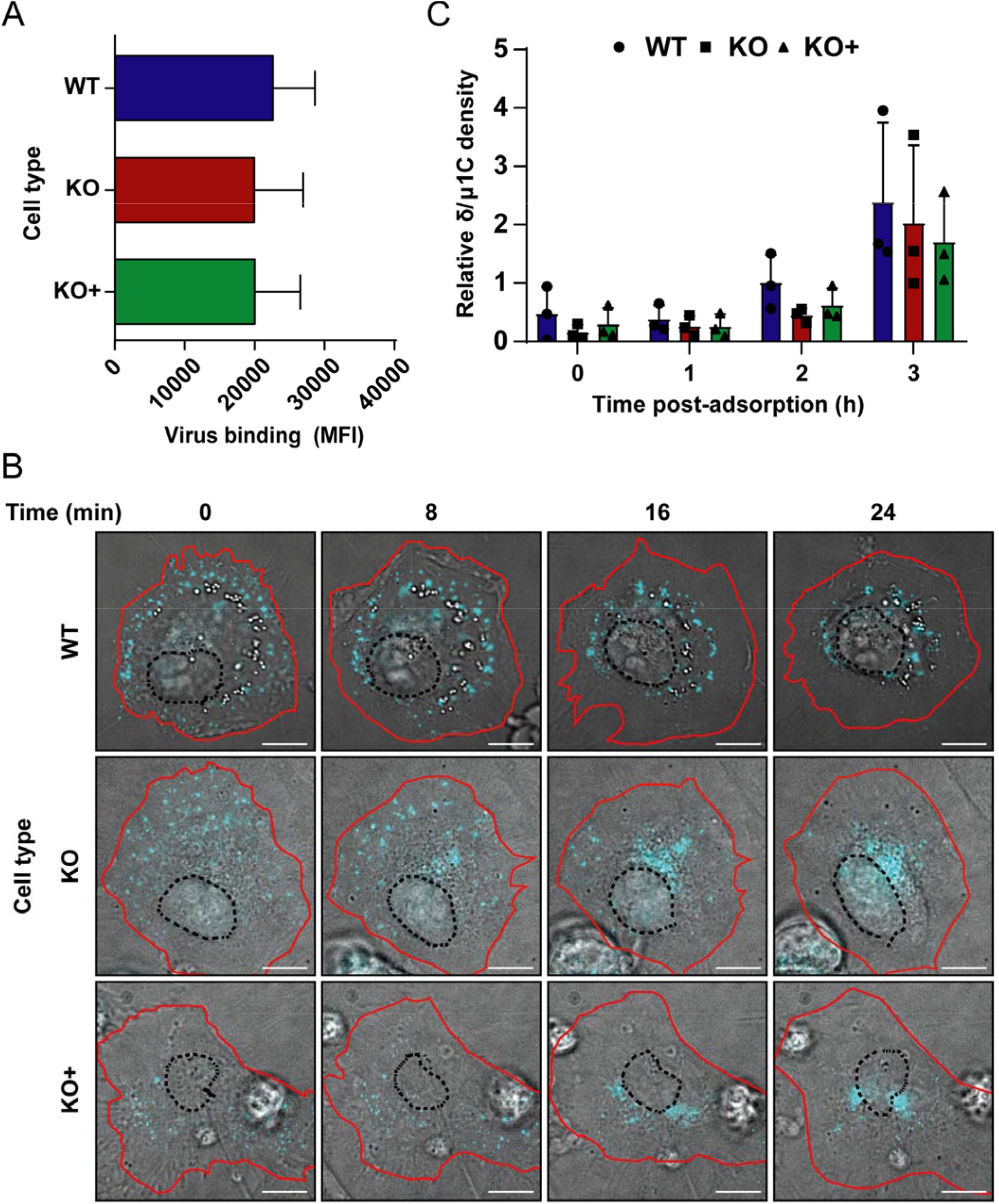
Binding, internalization, and uncoating are not disrupted by cholesterol accumulation in NPC1 KO HBMECs. (A) WT, KO, and KO+ HBMECs were adsorbed with Alexa 647 labeled-reovirus virions at an MOI of 10,000 particles/cell at 4°C for 1 h, fixed with 1% PFA, and analyzed for virus binding using flow cytometry. The results are presented as mean virus binding as determined by mean fluorescence intensity (MFI) of three independent experiments. Error bars indicated standard deviation. (B) WT, KO, and KO+ HBMECs were adsorbed with Alexa 647 labeled-reovirus virions at an MOI of 10,000 particles/cell at 4°C for 45 min and imaged using high magnification live-cell imaging, with images captured every ~ 25 seconds. Representative micrographs from videos at the indicated intervals are shown. Scale bars, 10 μm. (C) WT, KO, and KO+ HBMECs were adsorbed with reovirus virions at an MOI of 10,000 particles/cell at 4°C for 1 h and lysed at the intervals post-adsorption shown. Cell lysates were subjected to electrophoresis and immunoblotting using a reovirus-specific polyclonal rabbit antiserum. The results are presented as the mean ratio of the δ and μ1C bands from three independent experiments. Error bars indicate standard deviation. Differences are not significant, as determined by two-tailed unpaired t-test.

To determine whether NPC1 is required for reovirus to access the endocytic pathway of target cells, WT, KO, and KO+ cells were adsorbed with fluorescently-labeled reovirus particles and monitored for reovirus uptake using live-cell imaging. We found that the kinetics of reovirus internalization into WT, KO, and KO+ cells were comparable. High-magnification videos (Videos 1, 2, and 3) along with static images obtained at different intervals (Fig. 3B) demonstrate that attached reovirus particles internalize slowly in the first ~ 0 - 10 min post-adsorption. During this time, reovirus particles remain in the periphery, with a few particles coalescing to form large fluorescent puncta. Convergence of immunofluorescent signals suggests co-transport of multiple viral particles in the same endocytic compartment, similar to that observed during reovirus entry into neurons (44). After ~ 15 min post-adsorption, we observed rapid recruitment of almost every fluorescent puncta to the perinuclear region.

To more precisely define the movement of reovirus virions during entry, we analyzed the trajectories of individual fluorescent virions in Videos 1, 2, and 3 over 36 min using the Spot detector plugin function from Icy software. Trajectory colors change over time in which each color corresponds to an interval of ~ 7.5 min in the time-lapse videos (Videos 4, 5, and 6). Analysis of the time-dependent trajectories confirms observations made in the live-imaging videos. Thus, video-microscopic analysis demonstrates that reovirus virions are internalized rapidly into HBMECs and that virion uptake into the endocytic pathway is not impaired in the absence of NPC1.

Following internalization of reovirus virions, acid-dependent cathepsin proteases in late endosomes catalyze disassembly. During disassembly, proteolytic cleavage of the outermost capsid protein, σ3, exposes the membrane-penetration protein, μ1, which is subsequently cleaved to form a variety of intermediates that lead to penetration of the core particle into the cytoplasm (20–24, 27–30). Cells lacking NPC1 have increased endosomal pH and decreased cathepsin activity (45), which could impair reovirus uncoating. To determine whether NPC1 is required for reovirus disassembly, we defined the kinetics of reovirus outer-capsid proteolysis in WT, KO, and KO+ cells by following the formation of the δ cleavage fragment of the μ1 protein. Cells were adsorbed with reovirus virions, and viral proteins in cell lysates were visualized by immunoblotting at 0, 1, 2, and 3 h post-adsorption using a reovirus-specific antiserum. No significant differences in the kinetics of μ1 proteolysis were observed, with an initial δ cleavage product detected 2 h after adsorption in WT, KO, and KO+ cells (Fig. 3C). These data suggest that the cathepsins that catalyze reovirus disassembly are not impaired in NPC1 KO HBMECs. Collectively, these results demonstrate that NPC1 is not required for reovirus receptor binding, internalization, or disassembly.

### Escape of reovirus cores from endosomes is impaired in cells lacking NPC1

To determine whether NPC1 is required for escape of reovirus cores into the cytoplasm following disassembly in the endocytic compartment, we imaged cores in fixed cells by IF. Cells were adsorbed with fluorescently labeled reovirus virions and incubated in the presence of cycloheximide for 8 h post-adsorption to inhibit synthesis of new viral proteins and thus ensure detection of proteins from infecting viral particles. Cells were stained with a CD-63-specific antibody to label endosomes and an antiserum specific for reovirus cores and imaged using confocal microscopy. Small puncta consistent with reovirus cores were observed in WT and KO+ cells, while in KO cells, cores appeared to accumulate in larger puncta corresponding to endosomes (Fig. 4A). The distribution of virions, cores, and endosomes was determined to quantify the extent of colocalization. The results demonstrate frequent colocalization of cores and endosomes in KO cells (Manders coefficient [Mc]: ~ 0.7), while there was much less colocalization of cores and endosomes in WT and KO+ cells (Mc: ~ 0.3) (Fig. 4B). Colocalization of virions and cores also was more frequent in KO cells (Mc: ~ 0.45) than in WT (Mc: ~0.15) or KO+ (Mc: ~0.2) cells, whereas colocalization of virions and endosomes was comparable in all cell types (Mc: ~ 0.6). These data suggest that cores escape from endosomes more efficiently in the presence of NPC1.

**Fig. 4.**
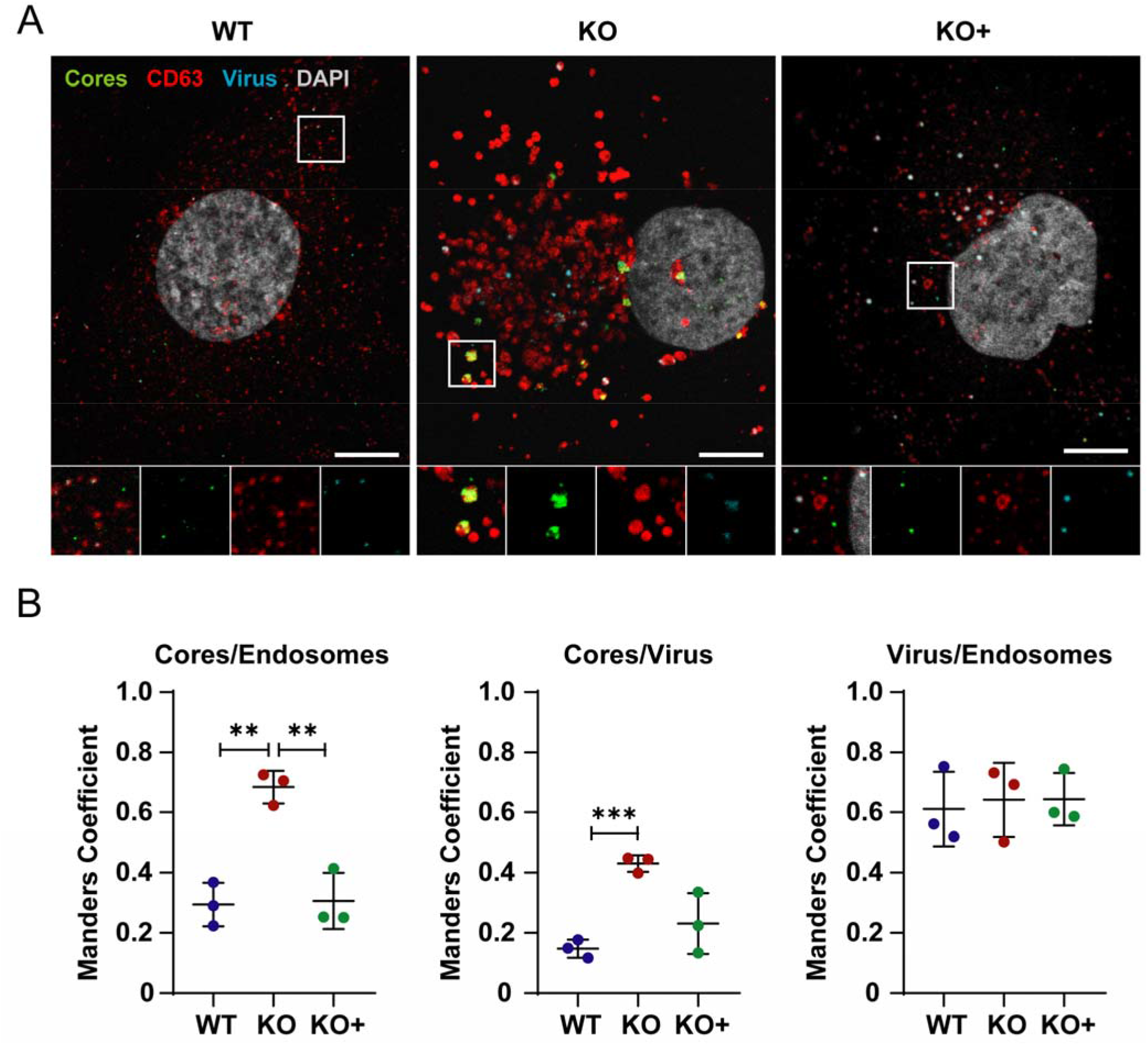
Cytosolic entry of reovirus cores. (A) WT, KO, and KO+ HBMECs were adsorbed with Alexa 647 labeled-reovirus virions at an MOI of 10,000 particles/cell at 37°C for 45 min and fixed with 4% PFA at 8 h post-adsorption. Cells were stained with DAPI, a CD-63-specific antibody to label endosomes, and an antiserum specific for reovirus cores, and imaged using confocal microscopy. Representative confocal micrographs are shown. (B) Colocalization of reovirus, cores, and endosomes was analyzed using the JaCoP plugin function from ImageJ. The results are presented as the mean colocalization (quantified by Manders coefficient) of ~ 50 cells from three independent experiments. Error bars indicate standard deviation. **, *P* < 0.01; ***, *P* < 0.001, as determined by two-tailed unpaired t-test.

To complement the imagining experiments, we quantified newly synthesized viral s4 mRNA using RT-qPCR. WT, KO, and KO+ cells were adsorbed with reovirus, RNA was isolated, and s4 transcripts were quantified at 0, 6, 12, and 24 h post-adsorption. We observed a statistically significant increase in total s4 RNA in WT and KO+ cells at 12 and 24 h post-adsorption relative to KO cells (Fig. 5). Together, these results suggest that NPC1 is required for release of transcriptionally active reovirus cores from endosomes into the cytoplasm.

**Fig. 5.**
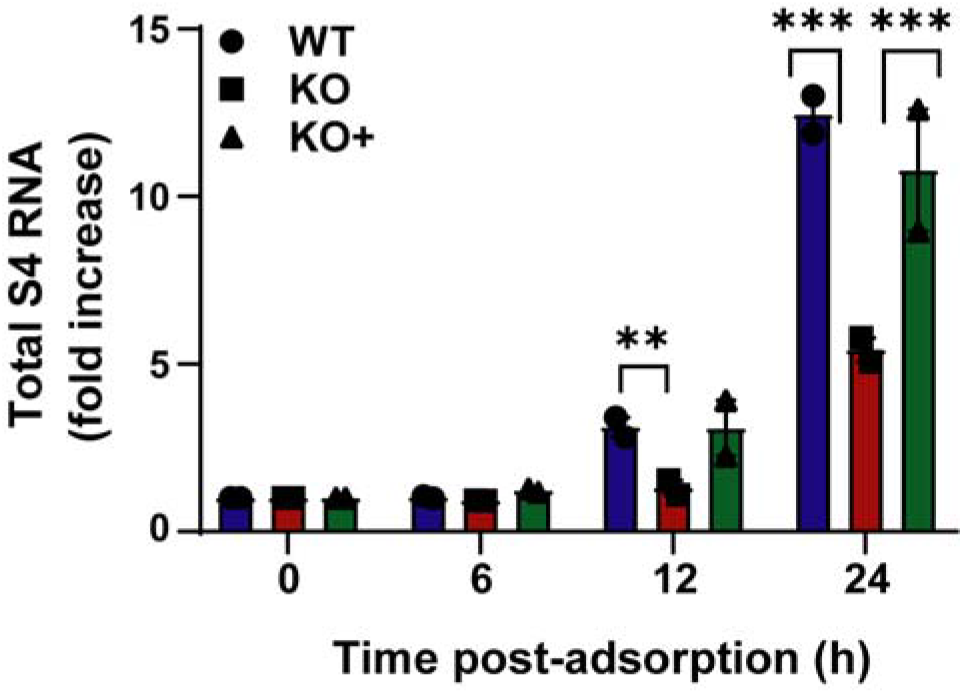
Synthesis of nascent RNA is reduced in NPC1 KO HBMECs. WT, KO, and KO+ HBMECs were adsorbed with reovirus virions at an MOI of 1 PFU/cell at 37°C for 1 h, lysed at the intervals post-adsorption shown, and assayed for positive-sense reovirus s4 RNA by RT-qPCR. The results are presented as the mean number of copies of reovirus s4 RNA by qPCR from two independent experiments. Error bars indicate standard errors of the mean. **, *P* < 0.01; ***, *P* < 0.001, as determined by t-test.

### Cholesterol homeostasis is required for reovirus entry

We thought it possible that NPC1 could serve as an endosomal receptor for reovirus and interact with one or more viral capsid proteins to enable core delivery into the cytoplasm, analogous to the function of NPC1 in Ebola virus infection (46, 47). Alternatively, NPC1 might be required to maintain an endosomal environment with appropriate cholesterol levels to allow cores to penetrate endosomes. To distinguish between these possibilities, we tested whether hydroxypropyl-β-cyclodextrin (HβCD), a cyclic oligosaccharide that triggers cholesterol release from the endo-lysosomal compartment (48, 49) and has been used to treat persons with Niemann-Pick disease type C (50, 51), for the capacity to overcome the effects of NPC1 deficiency on reovirus infection. To determine whether HβCD treatment redistributes cholesterol from endosomal membranes to a homogeneous distribution in the absence of NPC1, NPC1 KO HBMECs were treated with 1 mM HβCD, a non-toxic concentration (Fig. S3A), or PBS for 48 h prior to staining for the filipin III complex. Cells displaying cholesterol accumulation were distinguished from those with widely distributed cholesterol by quantifying the mean fluorescence intensity (MFI) of filipin III complex staining. Using this approach, an increase in MFI correlates with an increase in cholesterol accumulation. After HβCD treatment, KO cells displayed a significant redistribution of cholesterol, reducing its accumulation in endosomes and enhancing its distribution broadly throughout the cell, correlating with a statistically significant decrease in MFI (Fig. S3B,C). These data demonstrate that HβCD treatment promotes cholesterol efflux in KO cells, resulting in a cholesterol-distribution phenotype comparable to WT and KO+ cells (Fig. S3C).

Once we observed that HβCD treatment effectively redistributes cholesterol in KO cells and, thus, functionally complements NPC1 deficiency, we tested whether the reovirus entry defect in KO cells is due to the absence of NPC1 or impaired cholesterol homeostasis. WT, KO, and KO+ cells were pre-treated with 1 mM HβCD or PBS for 24 h, adsorbed with reovirus virions or ISVPs, and scored for reovirus infection by immunostaining. Remarkably, HβCD treatment rescued infection of KO cells by reovirus virions (Fig. 6) but did not appreciably affect infection of WT or KO+ cells. HβCD treatment also did not affect infection of WT, KO, or KO+ cells by ISVPs. These data demonstrate that endosomal cholesterol homeostasis regulates reovirus entry by enhancing penetration of reovirus core particles into the cytoplasm.

**Fig. 6.**
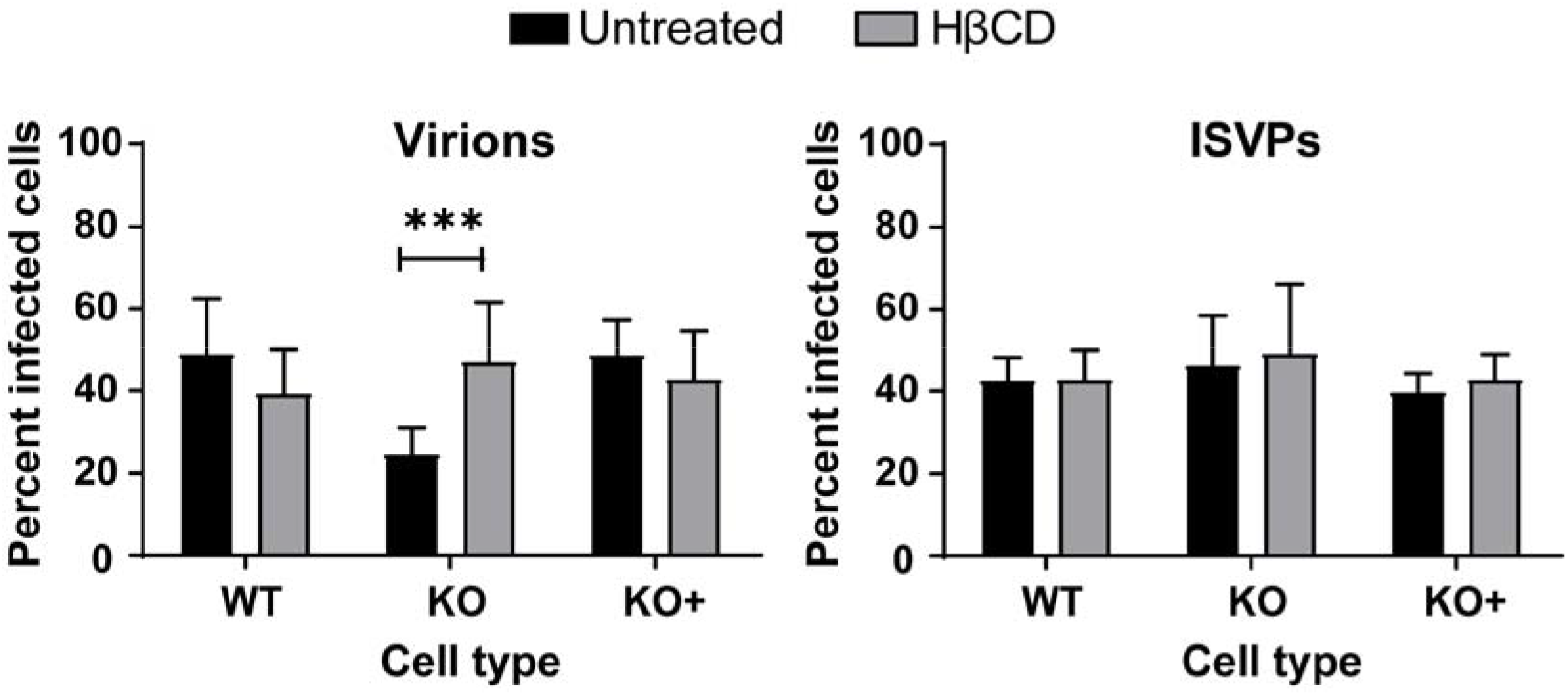
HβCD treatment restores reovirus infection of NPC1 KO HBMECs. WT, KO, and KO+ HBMECs were pretreated with 1 mM HβCD or PBS for 24 h, adsorbed with reovirus virions or ISVPs at MOIs of 10,000 or 100 particles/cell, respectively, and fixed at 18 h post-adsorption. The percentage of infected cells was determined by enumerating reovirus-infected cells following immunostaining with a reovirus-specific antiserum. The results are presented as the mean of three independent experiments. Error bars indicated standard deviation. ***, *P* < 0.001 as determined by two-tailed unpaired t-test.

## DISCUSSION

In this study, we identified NPC1 as a putative host factor required for reovirus infection using genome-wide CRISPR/Cas9 and siRNA-based cell-survival screens. NPC1 is an endolysosomal cholesterol transporter that mediates cholesterol homeostasis (32–34). Disruption of NPC1 results in cholesterol accumulation in late endosomes (Sup. Fig. 2C) and leads to Niemann-Pick disease type C, an autosomal-recessive neurodegenerative disorder (32). Early steps in reovirus infection, including receptor binding, acid-dependent proteolytic disassembly, and ISVP-to-ISVP* conversion have been well characterized (19). However, penetration of endosomal membranes and release of viral cores into the cytoplasm are poorly understood processes. We used CRISPR/Cas9 gene-targeted HBMECs lacking NPC1 expression to study the function of NPC1 in reovirus infection. We discovered that NPC1 is dispensable for viral binding to cell-surface receptors (Fig. 3A), internalization of viral particles (Fig. 3B), and disassembly of the viral outer capsid (Fig. 3C). However, NPC1 is required for efficient penetration of reovirus cores into the cytoplasm (Fig. 4). Treatment with HβCD reduces cholesterol accumulation in endosomes (Sup. Fig. 3B and 3C) and restores reovirus infectivity in NPC1 KO cells (Fig. 6). These findings suggest that regulation of cholesterol in endosomal compartments is essential for reovirus entry into host cells.

NPC1 is required for the replication of several enveloped viruses. The filoviruses Ebola virus and Marburg virus use NPC1 as an intracellular receptor (46, 47). NPC1 also functions in enveloped virus replication by maintaining cholesterol homeostasis. Disruption of cholesterol homeostasis by inhibiting NPC1 prevents entry and replication of dengue virus (52) and African swine fever virus (53) and impairs exosome-dependent release of hepatitis C virus (54). Additionally, NPC1 has been implicated in cell entry of quasi-enveloped forms of hepatitis A virus and hepatitis E virus (55, 56). However, NPC1 had not been previously appreciated to function in the replication of a nonenveloped virus.

We found that reovirus binding, internalization, and uncoating do not require NPC1, suggesting that NPC1 does not function as an intracellular receptor for reovirus. Instead, we found that cholesterol accumulation in the endocytic pathway diminishes the efficiency of reovirus core release into the cytoplasm. Using confocal microscopy, we visualized and quantified the distribution of fluoresceinated reovirus virions, reovirus cores, and late endosomes in infected cells (Fig. 4). Reovirus cores accumulate in the lumen of late endosomes in KO cells (Fig. 4A), while virions distribute to endosomes comparably in WT, KO, and KO+ (Fig. 4B). These findings suggest that cores do not escape from endosomes efficiently in the absence of NPC1. RNA synthesis, which occurs in the cytoplasm following release of cores from late endosomes, also was reduced in KO cells relative to WT and KO+ cells (Fig. 5), providing evidence that core escape from endosomes is required for initiation of transcription. It is not apparent how cholesterol accumulation in KO cells blocks core release from late endosomes.

In Niemann-Pick disease type C, disruption of cholesterol homeostasis causes changes in lipid composition of endosomal membranes (57, 58), inverting the ratio of phosphatidyl choline (PC) and phosphatidyl ethanolamine (PE). The change in PC:PE ratio may alter mechanical properties of endosomal membranes by inhibiting intra-endosomal membrane dynamics to favor negative curvature (57, 59). Membrane composition and dynamics can influence viral entry. Negative membrane curvature induced by addition of PE or the action of interferon-induced transmembrane protein 3 (IFITM3) impairs adenovirus protein VI-mediated membrane disruption (60) and enveloped virus fusion (61), respectively. Although reovirus virions are nonenveloped, entry of reovirus into cells also is inhibited by IFITM3 (62). Many nonenveloped viruses use membrane-modifying proteins with the capacity to interact, destabilize, and disrupt membranes to mediate genome release into the cytoplasm (12, 63). However, the role of specific lipids in these processes is not well defined.

During reovirus entry, ISVP-to-ISVP* conversion leads to release of myristoylated μ1N, which interacts with late endosomal membranes to facilitate release of cores into the cytoplasm (20–24). PE and PC concentrations in liposomes influence the efficiency of ISVP-to-ISVP* conversion (25). Therefore, it is possible that changes in membrane fluidity, width, or curvature caused by inversion of endosomal membrane PC:PE ratio in NPC1 KO cells impedes membrane insertion of μ1N or formation and expansion of the penetration pore. Additionally, accumulation of cholesterol within the endosomal compartment of NPC1 KO cells could limit recruitment of ISVP*s to membrane-inserted μ1N and the subsequent penetration of reovirus cores. Within the *Reoviridae* family, bluetongue virus (BTV) outer-capsid protein VP5 penetrates late endosomal membranes enriched in phospholipid lysobisphosphatidic acid (LBPA), which is dependent on the anionic charge and membrane fluidic properties of LBPA (64). LBPA-enriched late endosomes also are required for efficient rotavirus entry (65). Our data demonstrating the importance of cholesterol homeostasis in reovirus entry, along with the role of LBPA in BTV and rotavirus entry, suggest that the lipid composition of late endosomes influences nonenveloped virus entry and illuminate a potential new target for antiviral therapy.

Our findings parallel those of a companion study indicating a function for WD repeat-containing protein 81 (WDR81) in reovirus entry (66). WDR81 was identified in a CRISPR/Cas9 cell-survival screen using mouse embryo fibroblasts and found to be required for a step in reovirus entry that follows ISVP formation. WDR81 is required for the maturation of late endosomes by modulating levels of phosphatidylinositol 3-phosphate (67). These findings, coupled with our studies of NPC1, suggest that ISVPs formed in an altered endocytic compartment of cells lacking either WDR81 or NPC1 cannot launch replication, whereas ISVPs adsorbed to the surface of such cells can. We think that alterations in cholesterol distribution might govern this difference in ISVP behavior.

Cholesterol accumulation due to NPC1 dysfunction also can lead to alterations in the distribution of host proteins, such as annexin A2 (ANXA2), which was identified in our siRNA screen, and annexin A6 (ANXA6) (68). ANXA2 and ANXA6 are multifunctional proteins involved in endosomal trafficking, segregation of membrane lipids, and membrane curvature regulation through membrane-cytoskeleton rearrangements (69). Disruption of NPC1 leads to increased concentrations of ANXA2 and ANXA6 in late endosomes in response to cholesterol accumulation (70, 71). It is possible that cholesterol accumulation in cells lacking NPC1 similarly alters the distribution or function of WDR81. Thus, dysfunction of endosomal proteins in NPC1 KO cells might alter potential interactions of μ1N or the reovirus core with specific lipid microdomains or proteins and inhibit core release.

Genetic screens are useful approaches to identify host factors required for viral replication and provide valuable information about virus-cell interactions (72, 73). However, genetic screens frequently yield long lists of potential candidates, many of which are false-positives. To increase the likelihood of identifying host factors required for reovirus replication, we compared gene lists obtained from independent genome-wide CRISPR/Cas9 and siRNA-based cell-survival screens. Only 28 genes in the CRISPR/Cas9 screens using strains T1L and T3D were identified in the siRNA screen using strain T3SA+, 19 of which are ribosomal genes (Fig. 1B, C). Of the nine non-ribosomal genes, several encode proteins required for reovirus entry, including those involved in sialic acid biosynthesis and metabolism (*Nans* and *Neu*) (37, 38) and viral disassembly (*Ctsl*) (22).

Our findings indicate that NPC1, which was identified in both CRISPR/Cas9 and siRNA screens, is required for efficient release of reovirus cores into the cytoplasm by regulating cholesterol homeostasis. High-resolution studies showing the precise distribution of reovirus virions and cores within endosomes will be required to understand how NPC1 and cholesterol homeostasis regulate core release. These studies will allow us to answer the following new questions: Do cores interact with endosomal membranes in NPC1 KO cells? Does cholesterol impede interactions of cores with membranes? Are other lipids or proteins required for core release? Our ongoing work to answer these questions will clarify the functional elements of the reovirus entry pathway and lead to new approaches to block the entry of viruses that depend on tightly regulated cholesterol distribution in the endocytic pathway.

## MATERIALS AND METHODS

### Cells and viruses

HBMECs were cultured in growth medium (RPMI 1640 (Gibco) supplemented to contain 10% fetal bovine serum (FBS; VWR 97068-085), 10% Nu Serum (Corning), 1% MEM-vitamins (Corning), 1% sodium pyruvate (Gibco), 1% MEM non-essential amino acids (Gibco), 1% L-glutamine (Gibco), 1% penicillin/streptomycin (Gibco), and 0.1% amphotericin B (Sigma) or infection medium (growth medium containing 2% FBS). BV2 mouse microglial cells were cultured in BV2 maintenance medium (DMEM supplemented to contain 10% FBS, 1% penicillin/streptomycin, 1% sodium pyruvate, and 1% sodium bicarbonate) or selection medium (maintenance media supplemented with 4 μg/ml blasticidin (Thermo Fisher) and 2.5 μg/ml puromycin (Sigma-Aldrich)). HeLa cells were cultured in Dulbecco modified Eagle medium (Gibco) supplemented to contain 10% FBS, minimal essential medium nonessential amino acid solution (Gibco), 0.11 mg/mL of sodium pyruvate (Gibco), and 1% penicillin/ streptomycin, and 0.1% amphotericin B (Sigma). Spinner-adapted L929 cells (originally obtained from the Bernard Fields laboratory; ATCC CCL-1) were grown in either suspension or monolayers in Joklik’s modified Eagle’s minimal essential medium (US Biological; M3867) supplemented to contain 5% FBS, 2 mM L-glutamine, 100 units/ml penicillin, 100 μg/ml streptomycin, and 0.1% amphotericin B.

Reovirus strains T1L, T3D, T3SA+, and T1L M1-P208S, were prepared from laboratory stocks by plaque purification followed by 3 to 4 passages in L929 cells. T3SA+ contains nine genes from T1L and the S1 gene from T3C44-MA (36). T1L M1-P208S contains a point mutation in the M1 gene that causes viral factories to have a globular morphology similar to the morphology of factories formed by reovirus T3D (43) and can be readily scored for infection. Virions were purified from infected L929 cell lysates using cesium chloride gradient centrifugation as described (74). Viral titers were determined by plaque assay using L929 cells (75) and expressed as plaque forming units per ml (PFU/ml). Reovirus particle concentration was estimated by spectral absorbance of purified virions at 260 nm (optical density at 260 nm [OD_260_] of 1 = 2.1×10^12^ particles/ml) (76).

Fluorescent reovirus particles were prepared by diluting 6 × 10^12^ reovirus particles/ml in 50 mM sodium bicarbonate buffer and incubating with 20 μM Alexa Fluor™ 647 NHS Ester (Succinimidyl Ester) (Invitrogen, A37573) at room temperature (RT) for 90 min, protected from light (77). Labeled virions were dialyzed at 4°C overnight with 2-3 buffer exchanges to remove unreacted dye.

ISVPs were prepared by incubating 2 × 10^12^ purified reovirus particles with 200 μg/mL chymotrypsin (Sigma, C3142) at 37°C for 60 min (23). Digestion was terminated by the addition of PMSF to a final concentration of 2 mM. Virion-to-ISVP conversion was confirmed by SDS-PAGE and colloidal blue staining to assess the loss of σ3 and cleavage of μ1C to δ.

### Antibodies and dyes

Primary antibodies used for indirect immunofluorescence include anti-CD63 (1:250) (Thermofisher, #10628D), reovirus-specific polyclonal rabbit antiserum (1:1000) (78), and T1L core-specific rabbit antiserum (1:250) provided by Max Nibert (79). Alexa Fluor conjugated secondary antibodies (Thermo Fisher, #A11034, #A11030) were used to visualize antigen. Nuclei were stained with 4′,6-diamidino-2-phenylindole (DAPI, Invitrogen, D3571). Primary antibodies used for immunoblotting include reovirus-specific polyclonal rabbit antiserum, NPC1-specific polyclonal rabbit antiserum (Abcam, 134113), and mouse GAPDH monoclonal antibody for protein loading controls (Sigma, G8795). Anti-mouse IRDye680RD and anti-rabbit IRDye800CW (Licor) secondary antibodies were used.

### CRISPR Screen

The screen was conducted and transduction validated as described (80). BV2 cells were transduced with pXPR_101 lentivirus encoding Cas9 (Addgene; 52962) and propagated for 11 days with BV2 Maintenance Medium supplemented to contain blasticidin. These parental BV2 or BV2-Cas9 cells were transduced for 2 days with pXPR_011 expressing eGFP (Addgene; 59702) and a short guide RNA (sgRNA) targeting eGFP at a multiplicity of infection (MOI) of less than 1 PFU/cell. Cells were selected for 5 days with BV2 selection medium. The frequency of eGFP-expressing cells was quantified by flow cytometry.

The murine Asiago sgRNA CRISPR library contains six independent genome-wide pools, in which each pool contains unique sgRNAs targeting 20,077 mouse genes. Four pools of the Asiago library were transduced into 5 × 10^7^ BV2 cells at an MOI of 0.2 PFU/cell to establish four BV2 libraries. Two days post-transduction, cells were transferred to BV2 Selection Medium and propagated for 5 additional days. For each experimental condition, 10^7^ BV2 library cells expressing Cas9 and sgRNAs were seeded in duplicate into T175 tissue culture flasks (Greiner Bio-One). Cells were inoculated with Opti-MEM supplemented to contain PBS (mock) or reovirus strains T1L or T3D at an MOI of 100 PFU/cell. Cells were incubated at RT for 1 h, followed by the addition of 20 mL of DMEM supplemented to contain 10% FBS, 1% penicillin/streptomycin, 1% sodium pyruvate, and 1% sodium bicarbonate. After 2 days post-inoculation (dpi) (mock) or 9 dpi (T1L or T3D conditions), cells were harvested and genomic DNA (gDNA) was isolated from surviving cells using a QIAmp DNA Mini Kit (QIAGEN) according to the manufacturer’s instructions.

### CRISPR screen sequencing and analysis

Illumina sequencing and STARS analyses were conducted as described (81). The gDNA was aliquoted into a 96-well plate (Greiner Bio-One) with up to 10 μg gDNA in 50 μL of total volume per well. A polymerase chain reaction (PCR) master mix containing ExTaq DNA polymerase (Clontech), ExTaq buffer (Clontech), dNTPs, P5 stagger primer, and water was prepared. PCR master mix (40 μL) and 10 μL of a barcoded primer were added to each well containing gDNA. Samples were amplified using the following protocol: 95°C for 1 min, followed by 28 cycles of 94°C for 50 s, 52.5°C for 30 s, and 72°C for 30 s, and ending with a final 72°C extension for 10 min. PCR product was purified using Agencourt AMPure XP SPRI beads (Beckman Coulter) according to the manufacturer’s instructions. Samples were sequenced using a HiSeq 2000 (Illumina). Following deconvolution of the barcodes in the P7 primer, sgRNA sequences were mapped to a reference file of sgRNAs from the Asiago library. To account for the varying number of reads per condition, read counts per sgRNA were normalized to 10^7^ total reads per sample. Normalized values were then log-2 transformed. sgRNAs that were not detected were arbitrarily assigned a read count of 1. sgRNA frequencies were analyzed using STARS software to produce a rank ordered score for each gene. This score correlated with the sgRNA candidates that were above 10% of the total sequenced sgRNAs. Genes scoring above this threshold in either of the two independent subpools and in at least two of the four independent genome-wide pools were assigned a STAR score. In addition to the STAR score, screen results were compared using false discovery rate (FDR) analyses to monitor gene-specific signal versus background noise. Statistical values of independent replicates were averaged.

### Whole genome siRNA screen and analysis

The whole genome siRNA screen was conducted as described (35) using HeLa S3 cells and the Dharmacon ON-TARGETplus® SMARTpool® human siRNA library (Thermo Scientific) and strain T3SA+.

### Production of NPC1 KO and KO+ cell lines

HBMEC single-cell clones with ablation of the *NPC1* gene were engineered using CRISPR/Cas9-mediated gene editing as described (82) using an NPC1-specific gRNA (5′ GGCCTTGTCATTACTTGAGGGGG 3′, targeting nucleotides 768-790 of the human NPC1 mRNA). Single-cell clones were screened for the loss of NPC1 function by filipin III staining (82). Genotype of the selected NPC1 KO clones was confirmed by Sanger sequencing followed by amplification of the genomic DNA sequences flanking the gRNA target site using forward (5′ TCATAAACACACCAAACTTGGAATC 3′) and reverse (5′ TCCTGCGGCAGAGGTTTTC 3′) primers. Sequences of the NPC1 alleles were deconvoluted using CRISP-ID (83). To confirm the specificity of *Npc1* knockout, cells of a single clone were transduced with a retrovirus vector (pBabe-Puro) expressing human NPC1 as described (47).

### Indirect immunofluorescence staining

Cells were fixed with 4% paraformaldehyde (PFA, Electron Microscopy Sciences, 15712-s) in PBS^−/−^ at RT for 20 min, washed three times with PBS^−/−^, and permeabilized and blocked with 0.1% Triton X-100 and 2% FBS in PBS^−/−^ at RT for 20 min. Cells were incubated sequentially with primary antibody, Alexa Fluor-conjugated secondary antibody, and DAPI diluted in PBS^−/−^ containing 0.1% Triton X-100 and 2% FBS at RT for 30 to 60 min. For cholesterol labeling, fixed and permeabilized cells were incubated with 50 μg/ml filipin III (Sigma, SAE0088) diluted in PBS^−/−^ for 30 min. Coverslips were mounted using Prolong-gold (Molecular Probes). Confocal images were captured using a Leica-SP8 laser scanning confocal microscope equipped with an HCX PL APO 63X/1.4 N.A oil objective and processed using Fiji/ImageJ software.

### SDS-PAGE and Immunoblotting

Cells harvested for protein extraction were lysed in Radioimmunoprecipitation Assay buffer (*RIPA buffer*; Thermo Fisher) supplemented with 1X protease inhibitors (Thermo Fisher). Protein concentration was quantified by Bradford assay (Bio-Rad) following the manufacturer’s protocol. Samples for SDS-PAGE were diluted in 5X Laemmli sample buffer (Bio-Rad) containing 10% β-mercaptoethanol and incubated at 95°C for 10 min. Samples for detection of NPC1 were incubated at 70°C for 10 min to prevent aggregation. Equal amounts of protein were electrophoresed in 10% or 4-20% Mini-Protean TGX gels (Bio-Rad). Following electrophoresis, proteins were transferred to nitrocellulose membranes (Bio-Rad) for immunoblotting. Nitrocellulose membranes were incubated with 5% nonfat milk in TBS (50 mM Tris-HCl, pH 7.6; 150 mM NaCl) with 0.1% Tween 20 (TBS-T) and sequentially incubated with primary and secondary antibodies diluted in TBS-T at RT for 1 h. Immunoblot images were captured using an Odyssey CLx imaging system (Li-Cor) and protein bands were quantified using the Image Studio Lite software. Protein expression levels were normalized to GAPDH loading controls.

### Quantification of reovirus infectivity

In experiments comparing infectivity of reovirus in KO, KO+, and WT HBMECs, cells were adsorbed with 10,000 reovirus virions or 100 ISVPs diluted in Opti-MEM (Invitrogen) at 37°C for 1 h. Following adsorption, the inoculum was removed, and cells were incubated in infection medium for 18 h before fixing in ice-cold methanol. In experiments comparing reovirus infectivity in the presence or absence of HβCD, cells were treated with 1 mM HβCD or PBS for 24 h prior to adsorption with reovirus. Following adsorption, fresh 1 mM HβCD was added to the medium for 18 h before fixing in ice-cold methanol. Fixed cells were washed with PBS^−/−^, blocked with 1% bovine serum albumin (BSA), and incubated sequentially with reovirus-specific polyclonal rabbit antiserum, Alexa Fluor 488-conjugated anti-rabbit antibody, and DAPI in PBS^−/−^ containing 0.5% Triton X-100. Cells were imaged using a Lionheart FX automated imager (BioTek) equipped with a 20X air objective, taking four fields-of-view from duplicate samples. Images were processed and signals quantified using Gen5+ software (BioTek).

### Viral binding

KO, KO+, and WT HBMECs were detached from tissue-culture plates using CellStripper dissociation reagent (Corning), quenched with HBMEC medium, and washed with PBS^−/−^. Cells were resuspended in PBS^−/−^ at 10^6^ cells/ml and adsorbed with 10,000 Alexa Fluor 647-labeled reovirus virions/cell at 4°C for 1 h with agitation. After binding, cells were washed twice with PBS^−/−^ and fixed with 1% paraformaldehyde (PFA) supplemented with propidium iodide to determine cell viability. Cells were analyzed using an LSRII flow cytometer (BD Bioscience). Results were quantified using FlowJo V10 software.

### Live microscopy of reovirus internalization

KO, KO+, and WT HBMECs were plated on glass-bottom p35 plates and adsorbed with 10,000 Alexa 647-labeled reovirus virions/cell at 4°C for 45 min to synchronize infection. The inoculum was removed and replaced with fresh Opti-MEM without phenol-red medium supplemented with 2% FBS. Reovirus transport was imaged using a Leica DMI6000B fluorescence microscope with an HCX PL APO 63X/1.30 Gly objective. Fluorescence and brightfield images were collected from 0 to 36 min post adsorption every ~ 25 sec.

### Tracking of reovirus transport

Automated tracking of fluorescent reovirus particles in time-lapse images was conducted using Icy bioimage analysis software. Regions of interest (ROI) corresponding to the cell periphery were selected for tracking analysis using the Spot Detector plugin (84). The scale of the object (reovirus virions) to be analyzed was set at a size of ~7 pixels per spot, and the threshold sensitivity was set at 100. Parameters describing transport dynamics were considered as both diffusive and directed for running tracking analysis. Results are presented in colored time-dependent tracks.

### Quantification of reovirus cores

KO, KO+, and WT HBMECs were adsorbed with 10,000 Alexa Fluor 647-labeled reovirus virions at 37°C for 45 min. The inoculum was removed, and the cells were incubated in infection medium containing 100 μg/ml of cycloheximide for 8 h. After fixation, cells were permeabilized and stained with T1L core-specific rabbit polyclonal serum and anti-CD63 antibody. Confocal images were captured using a Leica-SP8 laser scanning confocal microscope equipped with an HCX PL APO 63X/1.4 N.A oil objective and processed using Fiji/ImageJ software. Colocalization of fluorescent reovirus virions (cyan puncta), reovirus cores (green puncta), and late endosomes (red puncta) was analyzed to differentiate infecting virions from cores released into the cytoplasm.

### RNA extraction and purification

Cells were lysed using TRIzol reagent (Invitrogen). RNA was extracted with chloroform and purified using a PureLink RNA minikit (Invitrogen) with DNase treatment according to the manufacturer’s instructions.

### S4 quantitative RT-PCR

Total S4 RNA was quantified using qScript XLT one-step RT-qPCR ToughMix, Low ROX (Quanta Bioscience) and T3D_S4_qPCR primers (Forward: GAAGCATTTGCCTCACCATAG, Reverse: GATCTGTCCAACTTGAGTGTATTG) according to the manufacturer’s instructions. The following RT-qPCR cycling protocol was used: cDNA synthesis (50°C for 10 min), initial denaturation (95°C for 1 min), and 40 PCR cycles (95°C for 10 s followed by a data collection step at 60°C for 1 min). S4 cDNA was detected using a fluorogenic probe (5′-FAM [fluorescent fluorescein]-AGCGCGCAAGAGGGATGGGA-BHQ [black hole quencher]-1-3′; Biosearch Technologies).

### Statistical analysis

All data were analyzed using Graphpad Prism 8. Figure legends specify the number of experimental repeats and the statistical test applied for each analysis. Differences were considered statistically significant when *P* values were less than 0.05.

## Acknowledgements

We thank members of the Dermody and Risco laboratories for many useful discussions and Dr. Pranav Danthi for review of the manuscript and sharing data from his laboratory prior to publication. We thank Dr. Martin Sachse for expert advice and review of the manuscript and Drs. Sylvia Gutiérrez-Erlandsson and Ana Oña for assistance with confocal microscopy. We are grateful to the UPMC Children’s Hospital of Pittsburgh Rangos Research Center Cell Imaging Core Laboratory for assistance with microscopy.

This work was supported in part by Public Health Service award R01 AI032539 (C.R. and T.S.D.) and the Heinz Endowments (T.S.D.) and grants BIO2015-68758-R and RTI2018-094445-B-100 from the Ministry of Science and Innovation of Spain (C.R.).

## Author contributions

**Conceived and designed experiments**: POG, GT, CR, and TSD. **Conducted experiments**: POG, GT, RKJ, BAM, RCO, and CBW. **Analyzed data**: POG and GT. **Contributed reagents/materials/analysis tools**: RKJ, RCO, CBW, and KC. **Wrote original draft**: POG, GT, and TSD. **Reviewed and edited paper**: POG, GT, RKJ, RT, IF, BAM, RCO, CBW, PAB, JS, KC, CR, and TSD.

## SUPPLEMENTAL MATERIALS

### FIGURES AND MOVIES

**Fig. S1. Effect on cholesterol distribution by disruption of NPC1 expression.** (A, B) Lysates of WT, KO, and KO+ HBMECs were subjected to electrophoresis and immunoblotting using an NPC1 antiserum. GAPDH was used as loading control. A representative immunoblot is shown. The results are presented as the mean of two independent experiments. Error bars indicate standard deviation. Statistical analysis was done by two-tailed unpaired t-test. (C) WT, KO, and KO+ HBMECs were stained with filipin III to detect cholesterol distribution. Representative images are shown. Scale bars, 10 μm. (D) WT, KO, and KO+ HBMECs were stained with filipin III and an anti-CD63 antibody to detect the subcellular localization of cholesterol. Representative images are shown. Scale bars, 10 μm.

**Fig. S2. Viral infectivity following adsorption by T1L, T3D, and T3SA+ virions.** (A, B) WT, KO, and KO+ HBMECs were adsorbed with reovirus virions at MOIs of 10,000 particles/cell, and fixed at 18 h post-adsorption. The percentage of infected cells was determined by enumerating reovirus-infected cells following immunostaining with a reovirus-specific antiserum. Error bars indicated standard deviation. **, *P* < 0.01; ***, *P* < 0.001, as determine by 2-way ANOVA, Tukey’s multiple comparisons test.

**Fig. S3. HβCD treatment restores cholesterol efflux in KO cells.** (A) WT, KO, and KO+ HBMECs were treated with HβCD at the concentrations shown for 48 h and assessed for viability using the Presto blue cell viability reagent. The results are presented as the mean cell viability of three independent experiments. Error bars indicated standard deviation. **, *P* < 0.01; ***, *P* < 0.001; ****, *P* < 0.0001, as determined by two-way ANOVA. (B, C) Cells were treated with 1 mM HβCD or PBS (mock) for 48 h, fixed with 4% PFA, stained with filipin III, and imaged using confocal microscopy. (B) The results are presented as the mean filipin III staining (quantified by MFI) of ~ 50 cells from three independent experiments. Error bars indicate the minimum and the maximum values. *, *P* < 0.05; ****, *P* < 0.0001, as determined by two-tailed unpaired t-test. (C) Representative images of cholesterol distribution in HβCD-treated and mock-treated cells are shown. Scale bars, 10 μm.

**VIDEO 1, 2, and 3. High-magnification, live-cell microscopy of fluorescent reovirus virion transport in WT, KO, and KO+ HBMECs.** (1) WT, (2) KO, and (3) KO+ cells were adsorbed with Alexa 647-labeled reovirus virions at an MOI of 10,000 particles/cell at 4°C for 45 min. Fluorescence and brightfield images were captured every ~ 25 seconds for 36 min.

**VIDEO 4, 5, and 6. Tracking of fluorescent reovirus virions recruited to a perinuclear region following entry.** Trajectories of reovirus virions during internalization into WT, KO, and KO+ HBMECs from videos 1, 2, and 3 were tracked with the spot-tracking plugin function of Icy-Bioimage analysis software (84). Cell contour was defined as a region of interest (ROI), and ~ 7 pixels/spot were monitored. The colored bar represents the trajectory depending on time, in which each color (from yellow to red) corresponds to an interval of ~ 7.5 min in the time-lapse videos. Scale bars, 10 μm.

