## Supplementary Fig 1 - 3 for "Reovirus infection is regulated by NPC1 and endosomal cholesterol homeostasis"

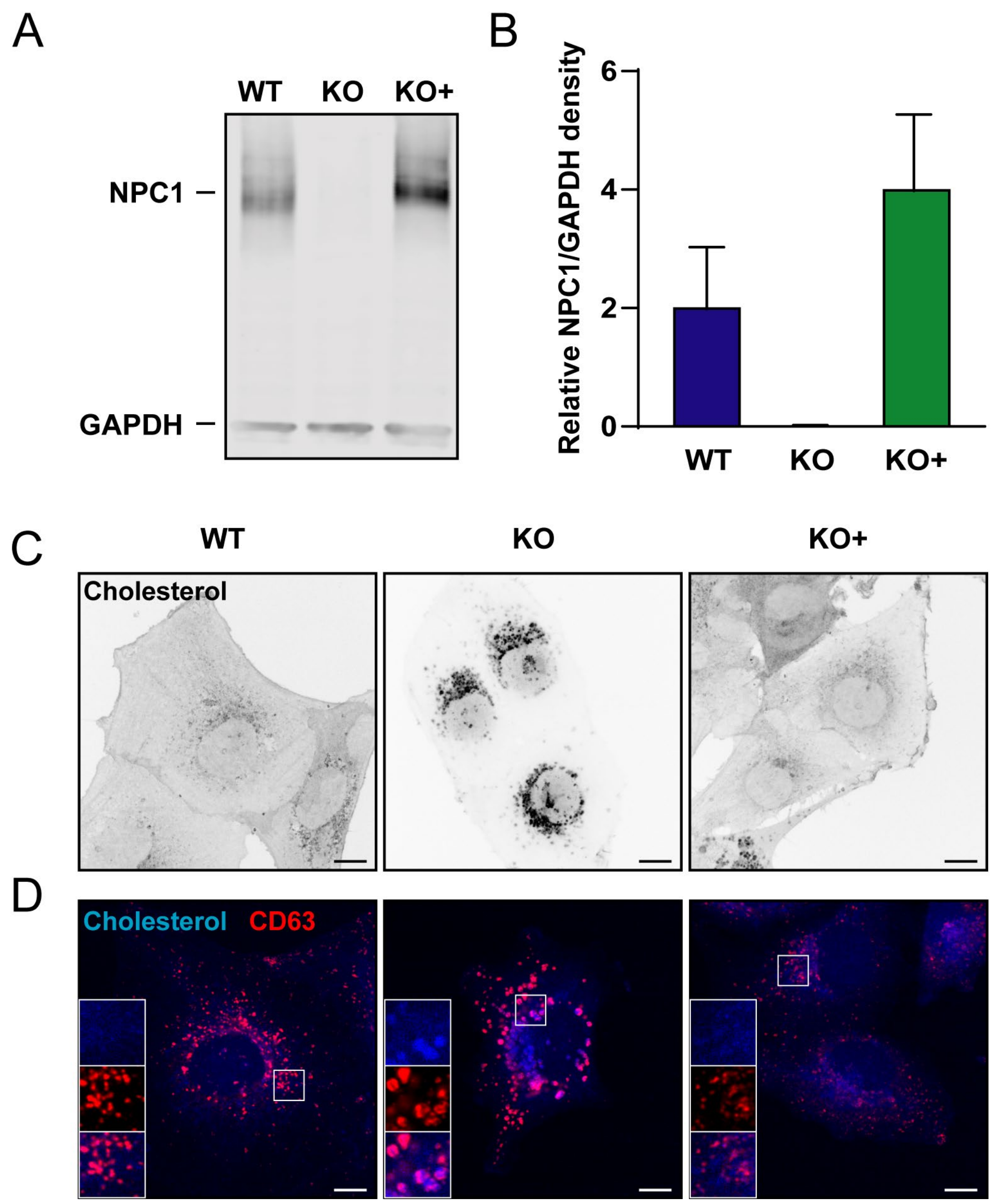


**FIG S1** Effect on cholesterol distribution by disruption of NPC1 expression. (A, B) Lysates of WT, KO, and KO+ HBMECs were subjected to electrophoresis and immunoblotting using an NPC1 antiserum. GAPDH was used as loading control. A representative immunoblot is shown. The results are presented as the mean of two independent experiments. Error bars indicate standard deviation. Statistical analysis was done by two-tailed unpaired t-test. (C) WT, KO, and KO+ HBMECs were stained with filipin III to detect cholesterol distribution. Representative images are shown. Scale bars, 10 µm. (D) WT, KO, and KO+ HBMECs were stained with filipin III and an anti-CD63 antibody to detect the subcellular localization of cholesterol. Representative images are shown. Scale bars, 10 µm.

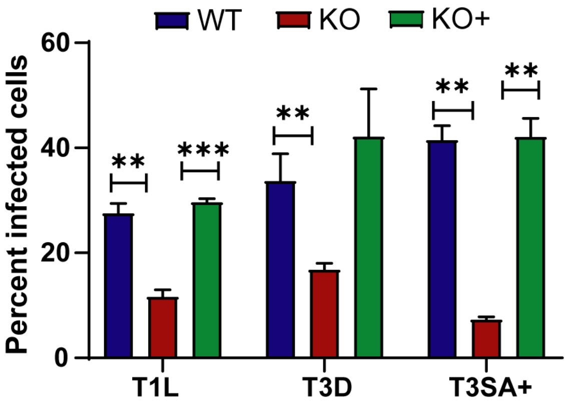


**FIG S2** Viral infectivity following adsorption by T1L, T3D, and T3SA+ virions. (A, B) WT, KO, and KO+ HBMECs were adsorbed with reovirus virions at MOIs of 10,000 particles/cell, and fixed at 18 h post-adsorption. The percentage of infected cells was determined by enumerating reovirus-infected cells following immunostaining with a reovirus-specific antiserum. Error bars indicated standard deviation. **, $P < 0.01$; ***, $P < 0.001$, as determine by 2 way ANOVA, Tukey's multiple comparisons test.

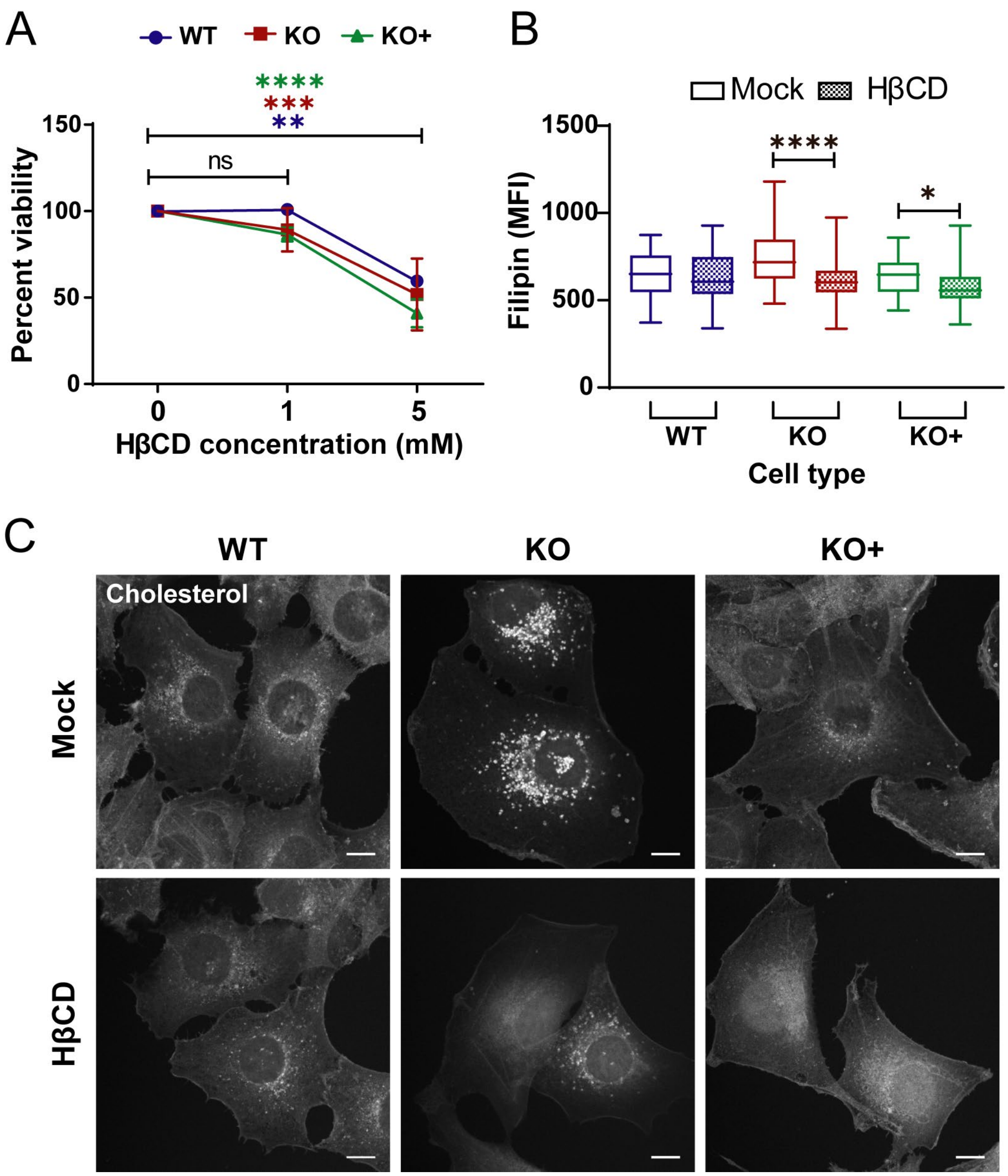


**FIG S3** HβCD treatment restores cholesterol efflux in KO cells. (A) WT, KO, and KO+ HBMECs were treated with HβCD at the concentrations shown for 48 h and assessed for viability using the Presto blue cell viability reagent. The results are presented as the mean cell viability of three independent experiments. Error bars indicated standard deviation. **, $P < 0.01$; ***, $P < 0.001$; ****, $P < 0.0001$, as determined by two-way ANOVA. (B, C) Cells were treated with 1 mM HβCD or PBS (mock) for 48 h, fixed with 4% PFA, stained with filipin III, and imaged using confocal microscopy. (B) The results are presented as the mean filipin III staining (quantified by MFI) of ~ 50 cells from three independent experiments. Error bars indicate the minimum and the maximum values. *, $P < 0.05$; ****, $P < 0.0001$, as determined by two-tailed unpaired t-test. (C) Representative images of cholesterol distribution in HβCD-treated and mock-treated cells are shown. Scale bars, 10 μm.
